# Engineering of an electrically charged hydrogel implanted into a traumatic brain injury model for stepwise neuronal tissue reconstruction

**DOI:** 10.1101/2022.02.16.480448

**Authors:** Satoshi Tanikawa, Yuki Ebisu, Tomáš Sedlačík, Shingo Semba, Takayuki Nonoyama, Akira Hirota, Taiga Takahashi, Kazushi Yamaguchi, Masamichi Imajo, Hinako Kato, Takuya Nishimura, Zen-ichi Tanei, Masumi Tsuda, Tomomi Nemoto, Jian Ping Gong, Shinya Tanaka

## Abstract

Neural regeneration is extremely difficult to achieve. In traumatic brain injuries, the loss of brain parenchyma volume hinders neural regeneration. In this study, neuronal tissue engineering was performed by using electrically charged hydrogels composed of cationic and anionic monomers in a 1:1 ratio (C1A1 hydrogel), which served as an effective scaffold for the attachment of neural stem cells (NSCs). In the 3D environment of porous C1A1 hydrogels engineered by the cryogelation technique, NSCs differentiated into neuroglial cells. The C1A1 porous hydrogel was implanted into brain defects in a mouse traumatic damage model. The VEGF-immersed C1A1 porous hydrogel promoted host-derived vascular network formation together with the infiltration of macrophages/microglia and astrocytes into the gel. Furthermore, the stepwise transplantation of GFP-labeled NSCs supported differentiation to glial and neuronal cells. Therefore, this two-step method for neural regeneration may become a new approach for therapeutic brain tissue reconstruction after brain damage in the future.

**One Sentence Summary:** Brain tissue reconstruction using charged hydrogel and stepwise NCS injection

## INTRODUCTION

The establishment of an efficient method for brain tissue regeneration is desired for the treatment of various diseases with brain parenchymal defects. Brain hemorrhage/infarction is a major cause of death and disability, with 13 million new cases per year^1,2^, and brain cancer, one of the most malignant human cancers, has 13.7 million new cases per year worldwide^3^. Both diseases are life-threatening and lead to enormous social and economic costs^2, 3^; thus, brain tissue regeneration has been studied^4^. However, there is no established method to date, and the development of an effective therapeutic strategy is urgently needed.

The brain is known to be vulnerable to ischemic damage, in which loss of the blood supply leads to the loss of neuroglial cells, the tissue framework, and extracellular matrix components, resulting in the loss of brain parenchyma volume and the formation of a cavity^1, 5^. This loss of tissue volume is irreversible, and spontaneous regeneration or wound healing processes do not occur, unlike in the skin, GI tract, and liver^6, 7^. One of the reasons is that the edge of the damaged brain tissue is sealed with a glial scar that prevents proper inflammatory cell migration and vascular extension for regeneration^8,9^. In general, neural regeneration is much more difficult in adults^10^ than in infants because the number of NSCs that are present as endogenous sources for neuroglial cells is low in adult brain tissue^11, 12, 13^.

Neural tissue engineering can solve these problems by using biomaterials to create a scaffold to compensate for volume loss, enabling cells to migrate into the damaged area and reconstruct the entire tissue structure^14–19^. Among various biomaterials, hydrogels have been shown to regulate the cellular differentiation of mesenchymal stem cells (MSCs)^20^, iPS cells^21^, and various cancer cells^22, 23^ and have recently been used for various biological applications^24^. Hydrogels are hydrophilic polymers commonly used in tissue engineering that can contain up to 90% water^25^ and have biological advantages, including a high oxygen content, nutrient permeability, and appropriate interfacial tension^26^. Natural substances have been used for hydrogels^19^, with various limitations, such as substantial variation^27^, the induction of immune reactions, and the release of components of tissue-derived pathogens^28^. To overcome these limitations, synthetic polymers have become widely used in hydrogels^19^. Neural tissue engineering using synthetic polymers and neurons may contribute to the regeneration of lost brain tissue^29^, but only a few reports have been published so far^5^.

In this study, to develop a new hydrogel material, we first focused on the electric charge of the substrate in cell adhesion^30–33^, and we found that an equal ratio of anionic and cationic monomers was most suitable for the attachment, growth, and differentiation of NSCs. Thereafter, the porous structure of the hydrogels was generated by cryogelation for 3D culture of neuroglial cells. In a traumatic brain injury model in mice, implantation of porous hydrogels with a similar stiffness to brain tissue induced host-derived neuroglial cell infiltration in the pores of the gels, along with vascularization. Furthermore, stepwise transplantation of NSCs into the hydrogels 2-3 weeks after implantation of the gels, where a vascular network formed and host glioneuronal cells had migrated, led to effective NSC survival, migration, and differentiation. These results indicate that porous hydrogels with a specific electric charge combined with stepwise NSC transplantation may represent a new therapeutic method for traumatic brain injuries.

## RESULTS

### Determination of the physical properties of hydrogels for culture of neural stem cells

To generate an optimum hydrogel for brain tissue reconstruction, we focused on the electric charge and investigated the conditions under which neural stem cells (NSCs) can efficiently attach and grow. Hydrogels with five different charges were synthesized using varying combinations of cationic and anionic monomers using 3-acryloylaminopropyl-trimethyl ammonium chloride (APTMA) and 2-acrylamido-2-methyl propane sulfonic acid, sodium salt (NaMPS) (Fig. 1a). The following five proportions were used: cation (C): anion (A) = 0:1, 1:5, 1:3, 1:1, and 3:1, which were designated C0A1, C1A5, C1A3, C1A1, and C3A1 gels, respectively. Noncharged dimethylacrylamide was used as a control. The surface electric charges, measured as the zeta potentials of the hydrogels, were as follows: C1A1, −245.9 ± 14.2 mV; C1A5, −113.6 ± 4.6 mV; C1A3, −76.3 ± 8.0 mV; C1A1, −5.9 ± 6.0 mV; C3A1, 100.0 ± 22.9 mV; and dimethylacrylamide, 2.1 ± 1.47 mV (Fig. 1b). Among these five hydrogels, we found that NSCs could efficiently attach and proliferate on the C1A1 gel (Fig. 1c). NSCs did not adhere to the negatively charged hydrogel, forming aggregates (C0A1, C1A5, and C1A3), while the NSCs temporarily attached to the positively charged hydrogels (C3A1), but they died over time. To investigate the mechanism of attachment, the involvement of surface proteins was examined^42^, and we found that the number of NSCs adhered to the C1A1 hydrogel was significantly decreased by treatment of the cells with a neutralizing antibody against integrin β1 (Supplementary Fig. 1).

**Fig 1.**
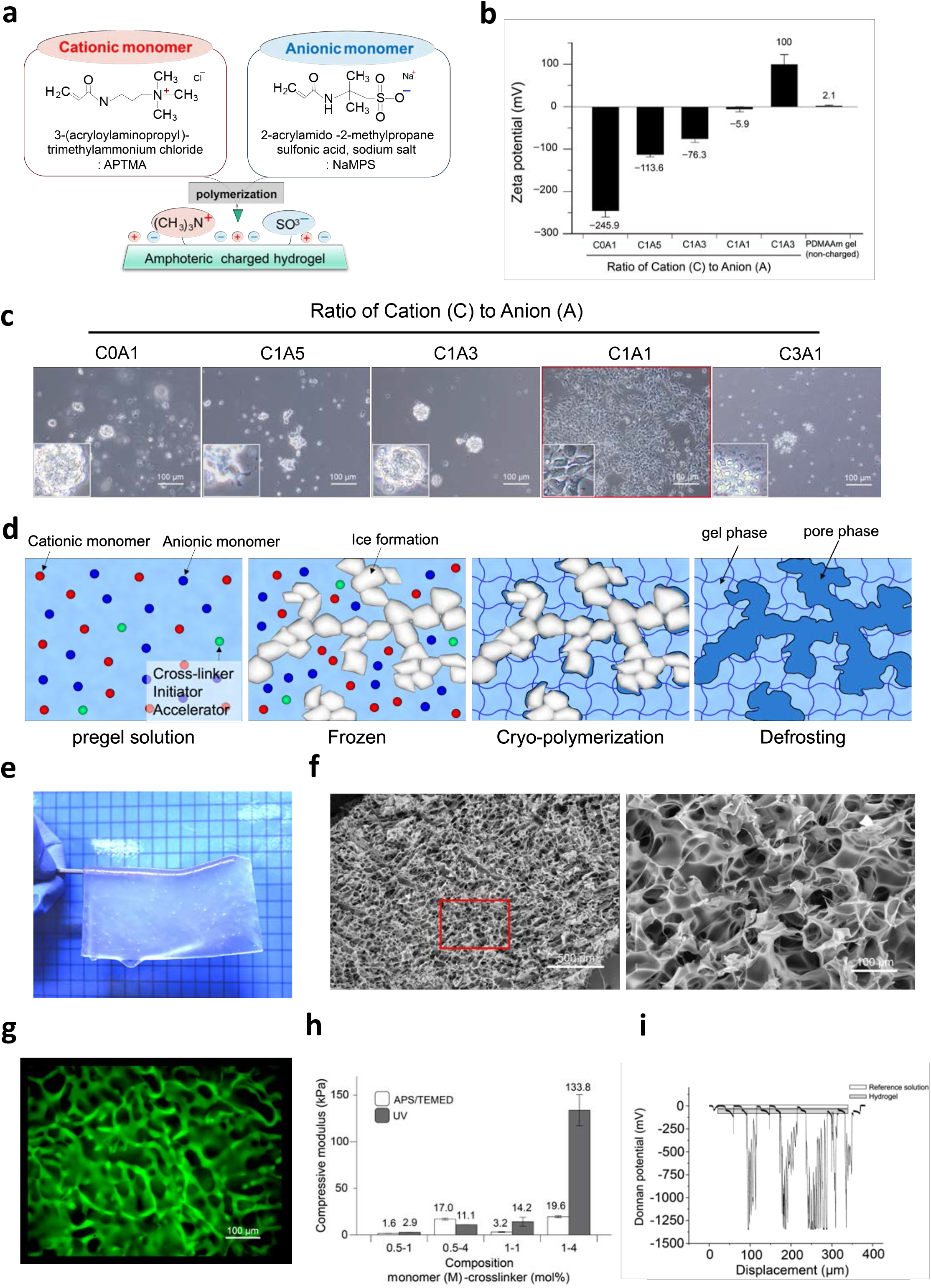
Engineering of an optimized hydrogel with a porous structure for 3D culture of NSCs. **a** and **b** Analysis of the various charged hydrogels for cellular attachment of NSCs. Schema of an amphoteric charged hydrogel. APTMA and NaMPS are shown as cationic monomers and anionic monomers, respectively (**a**). Zeta potential of each amphoteric hydrogel. A hydrogel synthesized with uncharged monomers (dimethylacrylamide) was used as a control (**b**). **c** Phasecontrast images of NSCs on differentially charged hydrogels such as C0A1, C1A5, C1A3, C1A1, and C3A1. NSCs were inoculated at a concentration of 1×10^5^/ml and cultured for 6 days with 10 ng/ml bFGF. **d** Schematic diagram of the cryogelation-formed porous hydrogel. The pregel solutions indicate the monomer solution of C1A1. Small circles in red, blue, and green represent APTMA, NaMPS and crosslinker, respectively (left panel). Solutions were frozen at −16°C overnight, and microsolid water structures with a white color were formed (2^nd^ left panel). To form the C1A1 hydrogel, polymerization at −16°C was performed as cryo-polymerization, and the formed network is shown as gray lines (2^nd^ right panel). After polymerization, the gels were warmed to defrost the samples, and a porous structure appeared (right panel). **e** Photograph of the semitransparent C1A1 porous hydrogel. **f** Scanning electron microscopy image of the porous C1A1 hydrogel. Interconnected macropores were observed. The area in the red box in the left panel is magnified on the right. White bars indicate 500 μm in the left panels and 100 μm in the right panels. **g** Fluorescence imaging of the C1A1 hydrogel incubated with fluorescein isothiocyanate (FITC). **h** Stiffness of the C1A1 hydrogels adjusted via the monomer and crosslinker concentrations. Open and closed boxes indicate APS/TEMED and UV cross linking, respectively. The average values and corresponding standard deviations were determined from at least three samples. **i** Measurement of the electric charge of the C1A1 porous hydrogel as the Donnan potential. White bar indicates the charge of the reference solution and the pore area (around 0 mV). Gray bar indicates the electric charge of surface to inside the porous gel.

For brain tissue reconstruction, a 3D culture system for NSCs is required; thus, a 3D cell culture substrate for C1A1 hydrogels was developed as a porous structure by using the cryogelation technique (Fig. 1d). Macroscopically, the C1A1 porous hydrogel was translucent (Fig. 1e). SEM demonstrated that interconnected pores with a maximum size of approximately 100 μm were formed with varying size (Fig. 1f and Supplementary Fig. 2a). Confocal microscopy using a fluorescent dye also delineated the interconnected pores of the C1A1 hydrogel (Fig. 1g, Supplementary movie 1).

To investigate the stiffness of the hydrogels in the context of the cellular microenvironment, eight types of C1A1 porous hydrogels with different stiffnesses were synthesized by altering the concentrations of monomer and crosslinker and employing two different polymerization methods: APS/TEMED or UV irradiation. These hydrogels with differential stiffnesses measured as moduli ranging from 1.6 to 133.8 kPa could be generated, and 0.5 M monomer and 1.0 mol% cross linker were found to provide moduli of approximately 1 kPa, simulating those of brain tissue (Fig. 1h). Lower monomer concentration achieved higher equilibrium water regain (*W*) (Supplementary Fig. 2b). Under this condition, the surface electric charge of the C1A1 porous hydrogel was confirmed to be neutral to slightly negative by measuring the Donnan potential (Fig. 1i). Therefore, the C1A1 porous hydrogel polymerized with APS/TEMED with 0.5 M monomer and 1.0 mol% cross linker was utilized for further experiments.

### Establishment of a traumatic brain injury model and therapeutic implantation of a C1A1 porous hydrogel into brain parenchymal defects

To evaluate the feasibility of using C1A1 porous hydrogels to compensate for brain defects *in vivo*, a traumatic brain injury model was established, and the C1A1 porous hydrogel was implanted into defects of the cerebral cortex in mice (Fig. 2a). For the formation of cylinder defects 1 mm in diameter and 1 mm in depth, the brain parenchyma was aspirated out using an aspirator equipped with a plastic loading tip 0.5 mm in diameter (Fig. 2b). The porous hydrogel was transplanted into the brain defect, and the condition of the gel was observed through the cranial window. Macroscopically, the transplanted gel seemed to become cloudy with time, suggesting regeneration of the meninges or infiltration of host cells into the porous hydrogel (Fig. 2c). The coronal section of the gel-transplanted brain exhibited a small amount of hemorrhage at the transplantation site on day 1, but no hemorrhage was observed after day 21. On day 56, the boundary between the brain parenchyma and the gels became obscured, while cavities clearly remained in the control brain (Fig. 2d). Microscopically, the C1A1 porous hydrogel was shown to be a light purple color with a porous structure by H&E staining (Fig. 2e). The volume of the defect was determined as a hypothetical cylinder. The section with the largest defect area in the coronal brain was selected, the area of the defect was measured, and the volume of the virtual cylinder was calculated by multiplying the measured area by the diameter of the defect. In the gel-implanted group, the volume of the cavity supported by the hydrogel was not changed, but in the control group, the diameter of defects expanded from day 21 (Fig. 2e, graph). In some cases, the hippocampus located under the defect protruded upward into the cavity without hydrogel. The porous structure of the C1A1 porous hydrogel was maintained for a long period of time (up to 294 days) after implantation (Supplementary Fig. 3a-b). The C1A1 porous hydrogel as a scaffold physically prevented the structural alterations associated with brain volume loss and maintained porous structures in the brain for a long period of time.

**Fig 2.**
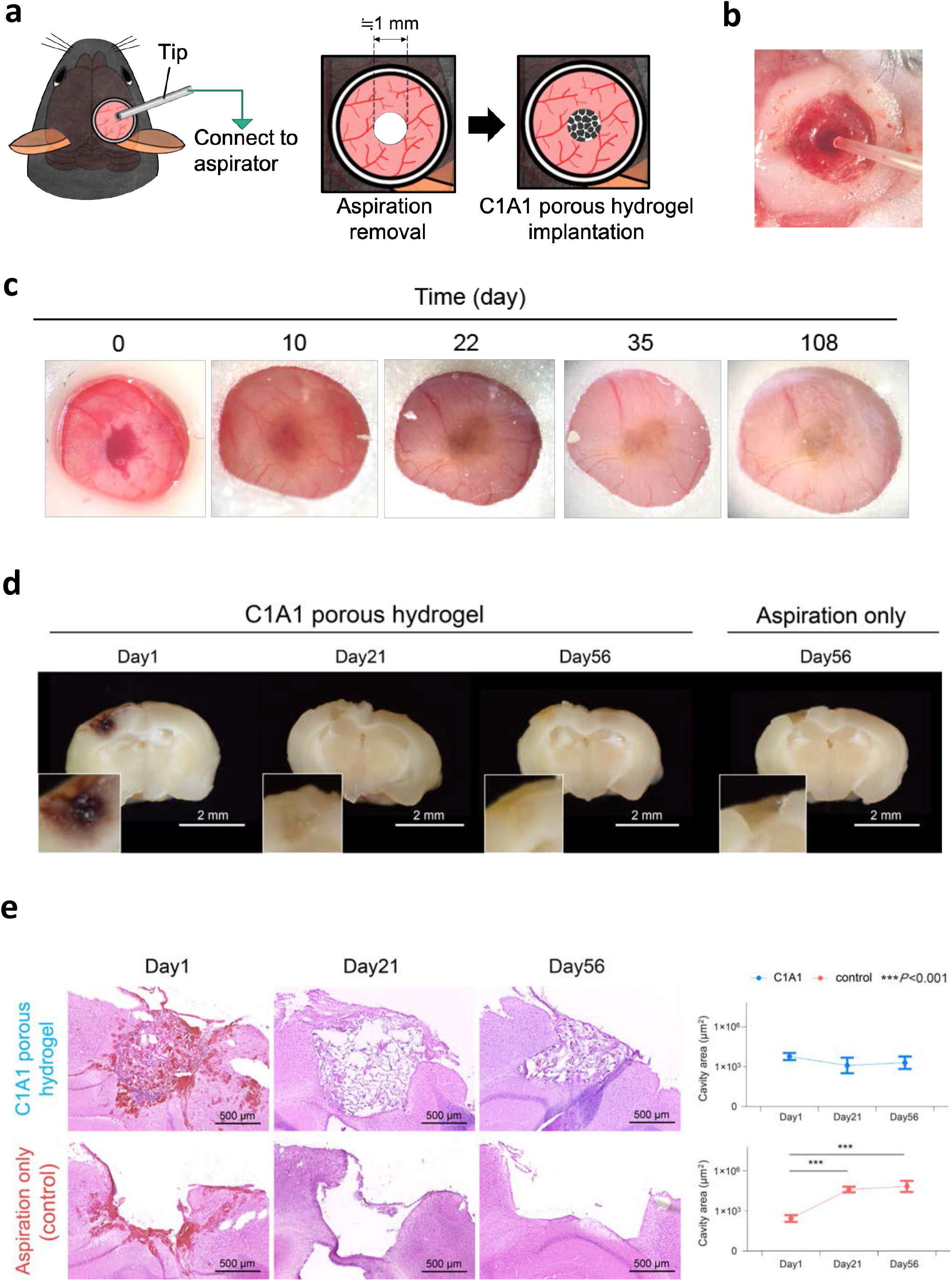
Establishment of a traumatic brain injury model and implantation of a C1A1 porous hydrogel into the mouse brain. **a** Illustration of the traumatic brain injury model in mice. The mouse brain was aspirated off as a cylinder shape, 1 mm in diameter and 1 mm in depth (left and middle), and a C1A1 porous hydrogel was implanted into the defect (right). **b** Stereoscopic image of a brain defect. **c** Stereomicroscopic analysis of the implanted C1A1 hydrogel in the brain defect through a cranial window on days 0, 10, 22, 35, and 108. **d** Macroscopic analysis of the coronal section of the implanted C1A1 porous hydrogel in the brain defect on days 1, 21, and 56. A brain defect without implantation is shown (right). Bars indicate 2 mm for the lower magnification in the photograph. White boxes indicate higher magnification images of the implanted gels. **e** Histological analysis of brain defects by H&E staining with (upper panels) or without (lower panels) implantation of the C1A1 porous hydrogel on days 1, 21, and 56 (upper panels). The volume of the defect was determined as a hypothetical cylinder. The section with the largest defect area in the coronal section was selected, and the area of the defect drawn from the cortical surface of the telencephalon to the hippocampus was measured with Fiji. The volume of the virtual cylinder was calculated by multiplying the measured area by the diameter of the defect. Each area was graphed as the mean ± SEM. \*\*\**P* < 0.001. n=5, each.

### Microscopic analysis of implanted C1A1 porous hydrogel in mouse brains

To investigate the potential of the host brain for tissue regeneration along with the implanted C1A1 porous hydrogel, immunofluorescence microscopy analysis was performed to evaluate the host cells penetrating or infiltrating the porous hydrogel. Various host-derived cells infiltrated into the implanted C1A1 porous hydrogel, and the total number of infiltrating cells measured by nuclear staining was significantly increased on day 21 and day 56 compared to day 1 (Fig. 3a, b). On day 1, most of the infiltrating cells were red blood cells. On day 21 and day 56, many immune cells and a small number of neurons positive for βIII tubulin were observed in the porous hydrogel (Fig. 3c-e). Neuronal axons were observed inside the pores that could be elongated from the host brain parenchymal tissues (Fig. 3d, circle). The numbers of infiltrating astrocytes also increased in a time-dependent manner, and by day 56, astrocytes rather than neurons could be predominantly observed in the porous hydrogel (Fig. 3f-h). Iba1-positive microglia were the most abundantly observed on day 21 and day 56 (Fig. 3i-j), and those cells engulfed brownish hemosiderin after day 21, suggesting phagocytosis of erythrocytes to resolve initial hemorrhage. Nestin-positive NSCs were also observed in the porous hydrogel (Fig. 3k), excluding the possibility of nestin-positive microglia in the infarcted area, as reported previously^43^.

**Fig 3.**
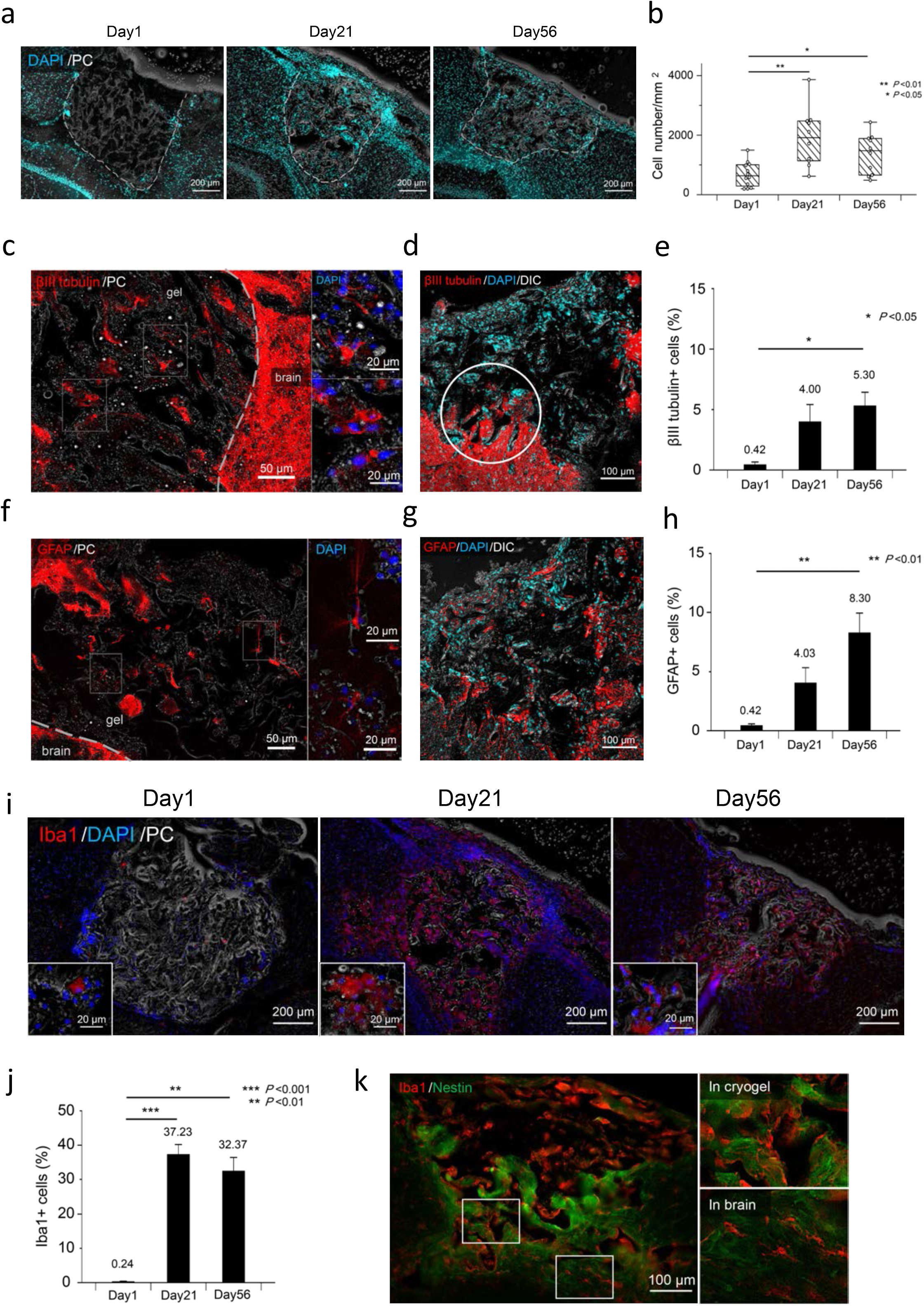
Fluorescence microscopy analysis of infiltration of host-derived cells into the implanted C1A1 porous hydrogel. **a** Analysis of total cell infiltration into the implanted hydrogel. Coronal sections of implanted hydrogels were analyzed by DAPI staining combined with phase contrast imaging (PC), and infiltrated cells in the C1A1 porous hydrogels were visualized on days 1, 21, and 56. **b** The numbers of cells were counted at each time point and are shown as box plots. \*\**P* < 0.01, \**P* < 0.05. n=5. **c**-**h** Analysis of neuronal cell (**c**-**e**) and glial cell (**f**-**h**) infiltration into the implanted hydrogel. Immunostaining of βIII tubulin (**c**) and GFAP **(f**) on day 21 (left panels). The dotted line indicates the border between the C1A1 hydrogel and the brain. Higher-magnification photographs of two areas in the left panels are displayed as two small panels on the right. Immunostaining of βIII tubulin (**d**) and GFAP (**g**) on day 56. The numbers of cells were counted at each time point and displayed as a bar graph using the mean ± SEM. \*\**P* < 0.01, \**P* < 0.05. n=5. **i** Analysis of macrophage/microglial infiltration into the implanted hydrogel. Immunostaining of Iba1 on days 1 (left panel), 21 (middle) and 56 (right) is displayed. The numbers of cells were counted at each time point and are shown as a bar graph using the mean ± SEM. \*\*\**P* < 0.001. \*\**P* < 0.01. **k** Immunostaining with Iba1 and nestin on day 56 (left panel). The right panels are an enlargement of the frame in the left panel.

It is worth analyzing the numbers of reactive astrocytes that may appear in brain parenchymal defects in general. The fluorescence intensity of GFAP around the cylindrical defects with or without hydrogel implantation was measured by dividing four tiered areas from the border of the hydrogel and brain parenchyma, designated 0-20, 20-70, 70-120, and 120-170 μm. As reported, a more than twofold increase in the number of activated astrocytes, termed gliosis, was observed in the control mice compared to the hydrogel-implanted mice in the area of 0-20 μm on day 21 (Supplementary Fig. 4a-b). On day 56, the numbers of reactive astrocytes in control mice were the same as those in hydrogel-implanted mice. Thus, on day 21, reactive astrocytes may migrate into the C1A1 porous hydrogel and prevent the formation of a clear glial scar at the boundary.

### VEGF promotes vascularization in the C1A1 porous hydrogel

Considering the effective brain tissue reconstruction in the implanted hydrogel, induction of the vascular network supplied from the host brain parenchyma should be important. Since transplantation of the C1A1 porous hydrogel alone failed to induce angiogenesis, the C1A1 porous hydrogel was immersed in VEGF, which promotes angiogenesis, and implanted into the mice brains. To confirm the formation of blood vessels in the implanted C1A1 porous hydrogel, *in vivo* live imaging using a two-photon microscope was performed, and vascularization was observed inside the gel on day 15 (Fig. 4a). Immunofluorescence on day 21 revealed infiltration of CD31-positive endothelial cells (Fig. 4b). Vascularization was also observed inside the pore in the long-term transplantation sample on day 200, and a-smooth muscle actin (αSMA) immunostaining revealed mature vascularization with the formation of a muscular layer in the vessel walls (Fig. 4c). Although the C1A1 porous hydrogel with VEGF induced the formation of a mature vascular network, the numbers of neuronal and glial cells were not significantly increased (data not shown).

**Fig 4.**
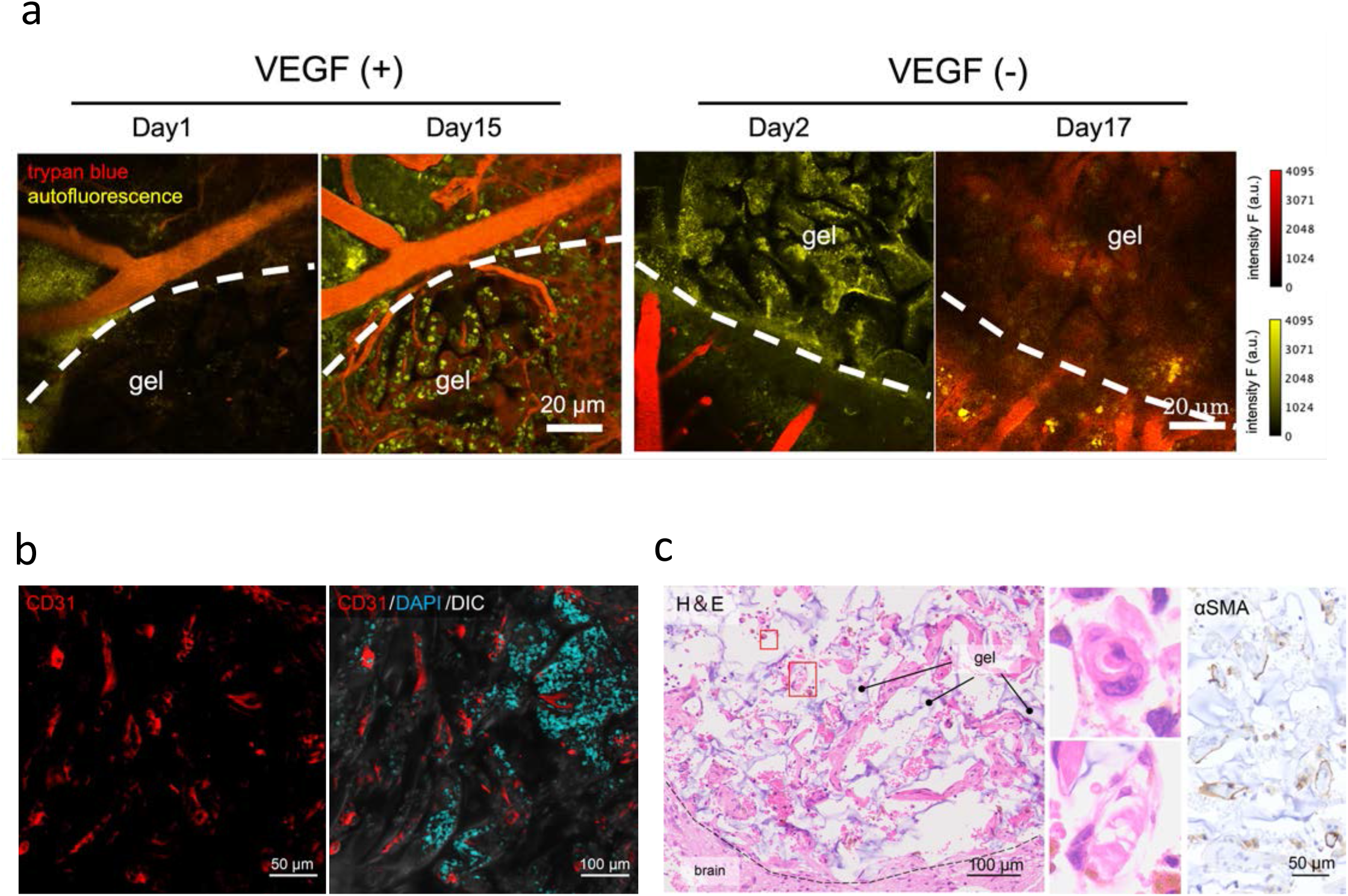
Vascular network formation in implanted VEGF-immersed C1A1 porous hydrogels. **a** *In vivo* live imaging of implanted C1A1 porous hydrogels by two-photon microscopy through a cranial window. C1A1 porous hydrogels were immersed in PBS with (left, n=3) and without (right, n=4) VEGF (50 ng/ml). In the VEGF (+) group, blood vessel formation was confirmed in two out of three mice, but in the VEGF (−) group, no blood vessels were formed in any of the four mice. Trypan blue was injected into the mice through the ocular vein, and the boundary areas were observed at the indicated time points. The dotted lines are the boundary between the C1A1 porous hydrogel and the brain tissue. Red indicates trypan blue (vessels), and yellow indicates cells that produce autofluorescence, such as immune cells. **b** Immunostaining of the implanted C1A1 porous hydrogel for CD31 on day 21 (left) in combination with DAPI (right). **c** Histological analysis of the implanted C1A1 porous hydrogel on day 200 (left panel). Light purple indicates the hydrogel. Higher-magnification photographs of two areas in the left panel with blood vessels are displayed as two small panels in the middle. Immunostaining for αSMA is shown (right).

### The C1A1 porous hydrogel acts as a scaffold for stepwise cell transplantation into brain defects following initial gel implantation

To obtain sufficient numbers of neurons, NSCs were transplanted into the C1A1 porous hydrogel. First, we examined the adhesiveness of NSCs *in vitro*. NSCs were seeded on the C1A1 porous hydrogel and cultured in medium supplemented with bFGF to maintain proliferation for one week. Subsequently, bFGF was removed, and the cells were incubated for one week to induce neuronal and glial differentiation. Neuronal cells were observed to be attached to C1A1 hydrogels (Fig. 5a, arrow). Neuronal differentiation and glial differentiation toward oligodendrocytes were visualized as positivity for βIII-tubulin and myelin basic protein (MBP), respectively (Fig. 5b, Supplementary movie 3). Several cells were confirmed to be positive for glial fibrous acidic protein (GFAP), a marker of astrocytes (data not shown). In some areas, neuronal processes appeared to be surrounded by MBP-positive oligodendrocyte processes (Fig. 5c, Supplementary movie 4). These data suggest that C1A1 porous hydrogels can be a scaffold for NSCs to grow and differentiate.

**Fig 5.**
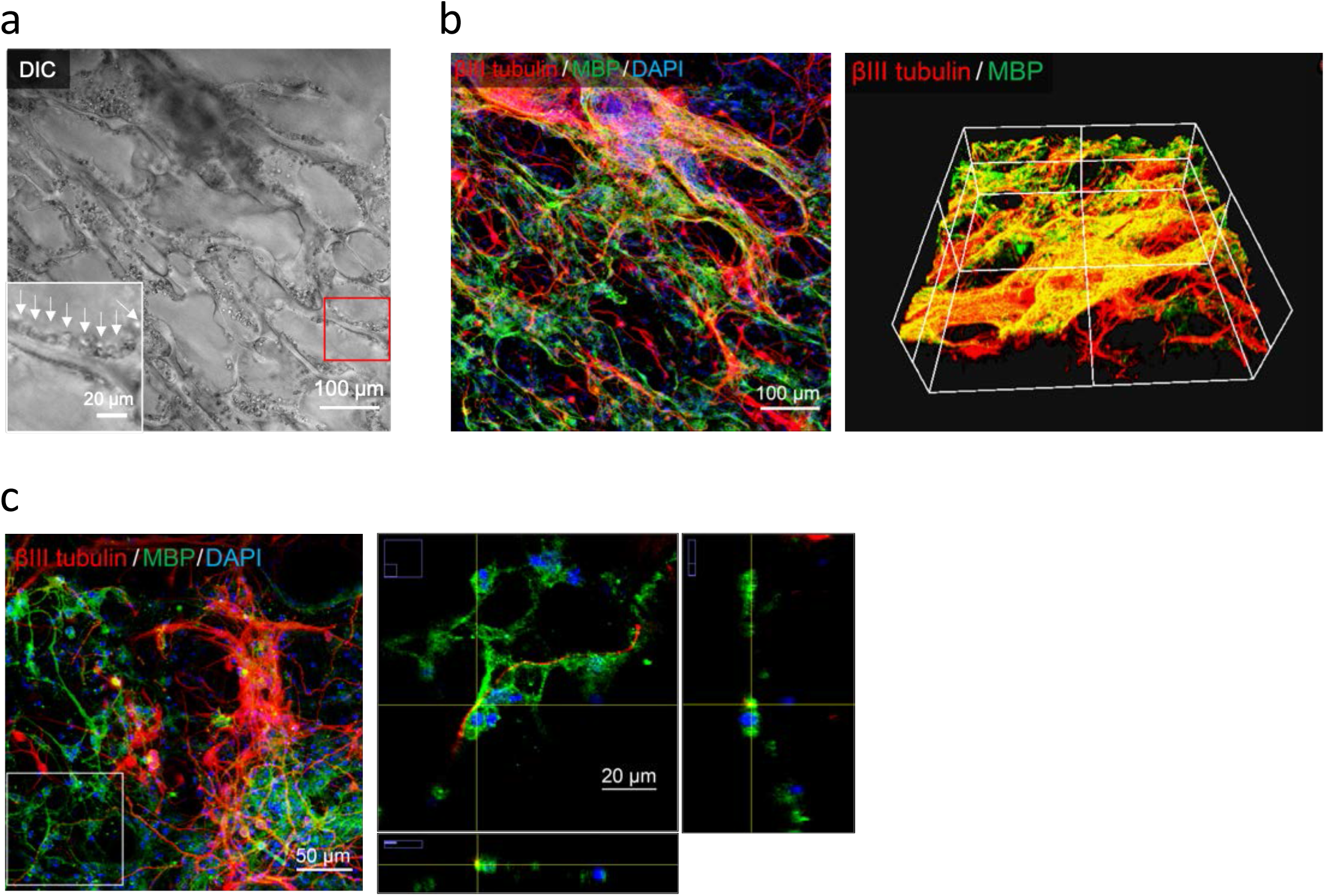
*In vitro* three-dimensional cultures of NSCs on the C1A1 porous hydrogel. **a** Differential interference contrast (DIC) image of NSCs 2 weeks after seeding into the C1A1 porous hydrogel. bFGF (10 ng/ml) was added daily for the first week, and bFGF was removed for the second week. The inset is a higher magnification of the area in the red square. **b** Immunofluorescence image. Three-dimensional cultures of neurons and glial cells were confirmed to stain red for βIII tubulin and green for myelin basic protein. Blue indicates nuclei visualized by DAPI staining. **c** Immunofluorescence images of areas of confirmed myelination. The white box indicated in the left panel is magnified and shown as the middle panel. The crossing point of the yellow vertical and horizontal lines is magnified and shown as lower and right panels. White bars indicate 50 μm and 20 μm in the left and middle panels, respectively.

NSCs labeled with GFP were injected into the implanted C1A1 porous hydrogel located in the mouse brain in a stepwise manner on day 21 after gel implantation when blood vessel formation was observed (Fig. 6a). On day 40 (64 days after gel transplantation), many cells could be observed in the gels (Figs. 6b and 6c). Fluorescence images demonstrated the presence of GFP-positive injected cells in transplanted C1A1 porous hydrogels; however, without porous hydrogel implantation, direct injection of NSCs alone or NSCs together with Matrigel did not support cell survival in the mouse brain (Fig. 6d-e). Some of the transplanted cells infiltrated the host brain tissues (Fig. 6f). Merging with differential interference contrast (DIC) images showed that host cells or transplanted cells could grow and migrate along the edge of the porous hydrogel (Fig. 6g). On day 40, most of the injected NSCs remained nestin positive due to stem cell properties, and some of the provided NSCs were labeled with GFAP as an astrocytic lineage, while small numbers of cells were positive for βIII-tubulin as differentiated neurons (Fig. 6h). After long-term observation on day 64, neuronal differentiation of the injected cells could be demonstrated in the porous hydrogels, together with the migration of neurons or the protrusion of neuronal axons into the host brain parenchyma (Fig. 6i). Similar results were obtained in NOD/Shi Jic-scid mice, with high survival of transplanted cells and vascularization (Fig. 6j).

**Fig 6.**
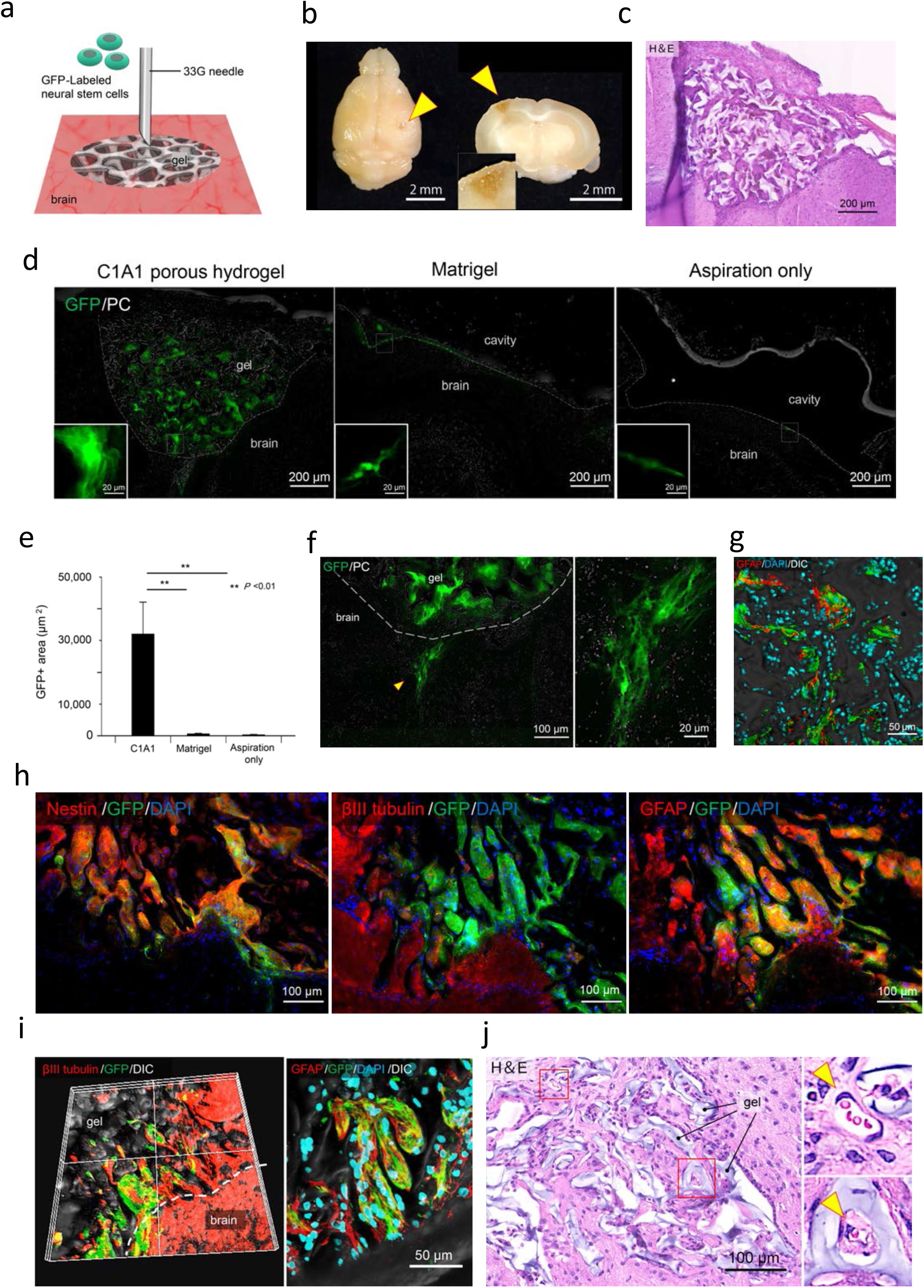
Stepwise transplantation of GFP-labeled NSCs into the implanted C1A1 porous hydrogel. **a** Diagram of NSC transplantation into the C1A1 porous hydrogel. Cell transplantation was performed approximately on day 21 after implantation of the C1A1 porous hydrogel (Table 1, right column). GFP-labeled NSCs were suspended in PBS at a concentration of 5×10^7^/ml. The needle was fixed at a depth of 1 mm from the surface of the hydrogel. The cell suspension was injected through a 33G needle in a volume of 2 μl, and the total number of injected cells was 100,000 cells. **b** Macroscopic photographs of NSC-transplanted brains. Yellow arrowheads indicate injected sites. The whole brain (left) and a coronal section (right) of the implanted C1A1 porous hydrogel at a higher magnification (inset of right). **c** Histological analysis of the transplanted site by H&E staining. **d** Detection of GFP-labeled NSCs transplanted into the C1A1 porous hydrogel by fluorescence microscopy (left). As a control, NSCs suspended in Matrigel (middle) or PBS alone (right) were transplanted. The insets show a higher magnification view of GFP-positive cells. **e** The GFP-positive areas were measured and displayed as a bar graph with the mean ± SEM. \*\**P* < 0.01. **f** Fluorescence microscopy of transplanted NSCs migrating into the host brain parenchyma (left). The dotted line indicates the boundary between the C1A1 porous hydrogel and the brain. The yellow arrowhead indicates the migrated area, and a higher magnification is shown (right). **g** Fluorescence microscopy of GFP-NSCs and host-derived GFAP-positive glial cells (red) migrating along the edge of the C1A1 porous hydrogel. **h** Fluorescence microscopy of differentiated transplanted GFP-NSCs on day 40. Immunofluorescence using antibodies against nestin (left), βIII tubulin (middle), and GFAP (right) is shown. **i** Fluorescence microscopy of infiltration of host neural (left) and glial cells (right) on day 64 after cell transplantation. **j** Histological analysis of NSCs transplanted into the C1A1 porous hydrogel on day 36 after cell transplantation in NOD/Shi Jic-scid mice (by H&E staining). Higher-magnification photographs of the red boxes are displayed as small panels on the right. Arrowheads indicate blood vessels.

## DISCUSSION

In this study, we demonstrated that C1A1 electrically charged porous hydrogels can serve as scaffolds for brain parenchymal defects, and stepwise transplantation of NSCs into the hydrogel following gel implantation may induce the reconstruction of brain tissue along with the implanted hydrogels. Currently, various biomaterials have been used in human regenerative medicine, and biomedical engineering for specific tissues has become an important method to compensate for organ dysfunction.

To create the ideal substrate for brain tissue reconstruction, hydrogels were employed because the stiffness of hydrogels can be adjusted to that of human brain tissue. It is well known that the stiffness of the cell culture substrate can regulate the direction of differentiation of MSCs, with a soft gel inducing neuronal cells and a hard gel inducing chondrocytes^20^. Previously, we reported that negatively charged PNaMPS gel has the potential to promote the differentiation of MSCs into chondrocytes for articular regeneration^44^. It should be noted that we first used PNaMPS gels, which were successful in culturing MSCs but failed in culturing NSCs. Thus, we focused on the electric charge of the hydrogels and discovered that the combination of cations and anion monomers at a 1:1 ratio meant that the C1A1 hydrogel can function as a scaffold for NSCs to attach and grow, with differentiation potential.

The surface potential of the substrate induces the adsorption of various proteins and has been reported to be involved in cell adhesion^45, 32^. The C1A1 hydrogel adhered well to NSCs but less well to MSCs and various cancer cell lines (data not shown). It will be interesting to clarify the mechanism by which NSCs efficiently adhered to the C1A1 hydrogels. Thus far, the involvement of integrin β1 has been suggested (Supplementary Figs. 1a, 1b). Recently, we and others reported that several molecules, including integrins or the ion channel family of proteins, may function as mechanoreceptors in fibroblasts or cancer cells^22, 46^, and further study is required to uncover the precise mechanism of specific interactions between hydrogel charge and cell surface proteins of NSCs.

To investigate the feasibility of using C1A1 porous hydrogels for brain tissue reconstruction, we established a brain damage model in mice, and then porous hydrogels were implanted. In the process of classical wound healing, four sequential modes comprising hemostasis and thrombus formation, inflammatory cell infiltration, granulation tissue formation, and remodeling are proposed^47^. In brain damage, inadequate production of extracellular matrix as a scaffold by astrocytes prevents granulation, including vascular formation, leading to a lack of neuronal repair^6^. In our model, initial brain defects led to bleeding and inflammation, and granulation tissue-like structures were formed in the group implanted with the C1A1 porous hydrogel, which functioned as the scaffold for cellular migration and growth, compensating for the insufficient production of extracellular matrix.

Recent studies have reported that inflammation is required for brain regeneration^48^ and is involved in axonal regeneration in the central nervous system^49^. In fact, the infiltration of Iba1-positive macrophages/microglia was prominent on day 21 rather than at later stages, and the numbers of neurons and glial cells were more pronounced in highly inflamed specimens (data not shown). In addition, angiogenesis into the gel is critical for efficient brain tissue reconstruction, and we succeeded in the formation of vascular networks in porous hydrogels by adding VEGF. Inflammatory cells such as macrophages/microglia are important for angiogenesis^50, 51^. Furthermore, hydrogels are used as drug delivery systems^19^; in fact, we have reported that NaMPS gel has a reservoir function^52^. Thus, the reservoir function of C1A1 porous hydrogels for endogenously secreted growth factors may be involved in angiogenesis, leading to neuronal tissue reconstruction.

The stepwise transplantation of GFP-labeled iPS-derived NSCs achieved efficient reconstruction of brain tissue. We achieved a high engraftment rate of transplanted cells after 2-3 weeks. The survival rate of transplanted cells has been reported to be low owing to the harsh environment that lacks extracellular matrix^53^. However, the viability of transplanted cells in this study was high, probably because the porous scaffold supported vascular network formation at the appropriate time, as confirmed by *in vivo* live imaging. Histologically, many inflammatory cells were still present 2-3 weeks after transplantation (Fig. 3e). Whether inflammatory cells have a positive or negative effect on transplanted cells seems to be controversial, but some reports indicate that inflammation has a supportive effect on tissue regeneration^48, 49, 54^. The inflammatory process and the best timing of transplantation for survival and differentiation of NSCs should be further investigated. Experiments using the hydrogel *in vivo* showed that the C1A1 porous hydrogel is covalently polymerized and basically has no biodegradability. Currently, the absence of biodegradability of biomaterials used for the preservation of brain function is controversial. To date, we have not observed any behavioral abnormalities or deaths in any of the mice implanted with the gels. Further study is required to determine the balance between appropriate biodegradability and adverse effects.

In this study, we established a basic strategy for brain tissue reconstruction using newly generated novel hydrogels and transplantation of NSCs in a stepwise manner. We hope that this approach will become a standard therapeutic procedure in the future.

## MATERIALS AND METHODS

### Preparation of hydrogels with various charges

Hydrogel sheets with a thickness of 1 mm were synthesized by using two parallel glass plates. To synthesize the differentially charged hydrogel, various combinations of a cationic monomer, 3-(acryloylaminopropyl)-trimethylammonium chloride (APTMA, Tokyo Chemical Industry Co., Ltd., Tokyo, Japan), and an anionic monomer, 2-acrylamido-2-methylpropane sulfonic acid, sodium salt (NaMPS, Sigma-Aldrich, St. Louis, MO, USA), were used. Dimethylacrylamide (Sigma-Aldrich) was employed as a neutral monomer. Five differentially charged hydrogels were constructed by using ratios of cationic (C) and anionic (A) monomers of 0:1, 1:5, 1:3, 1:1, and 3:1, which were designated C0A1, C1A5, C1A3, C1A1, and C3A1 gels, respectively. Polymerization of 1 M monomer was performed with 4 mol% N,N’-methylene-bis-acrylamide (Sigma-Aldrich) as a crosslinker with 0.1% (v/v) N,N,N’,N’-tetramethylethylenediamine (TEMED, Wako Pure Chemical Ind., Ltd., Japan) and 0.4% (v/v) ammonium persulfate (APS, Wako) at room temperature overnight. Alternatively, UV irradiation was used for polymerization. Polymerized hydrogels were immersed in a large amount of phosphate-buffered saline (PBS) to remove residual unpolymerized chemicals for 1 week, and the PBS was refreshed every day. Subsequently, gel disks were punched out of the gel sheet with a hole punch and sterilized in an autoclave at 120°C for 20 minutes. The gel disks were then placed on polystyrene tissue culture dishes and equilibrated in cell culture medium overnight.

#### Preparation of the C1A1 porous hydrogel via cryogelation

For the preparation of porous hydrogels, cryogelation was employed using C1A1 hydrogels. The aqueous solution containing monomers, crosslinker, 4 mg/mL TEMED, and 0.5 mg/mL APS was purged with argon for 20 minutes to remove dissolved oxygen, inhibiting radical polymerization. The solution was cooled in an ice water bath to slow the speed of polymerization. The glass molds precooled in a cooling bath (−16°C) and injected with reaction solution were inserted into a zipper plastic bag and immediately placed in an ethanol cooling bath (NCB-3300, TOKYO RIKAKIKAI Co, Ltd., Tokyo, Japan). Frozen samples were kept at −16°C overnight to allow crosslinking polymerization. Samples with a UV initiator (0.5 mg/mL alpha-keto, Wako) were irradiated. After polymerization, the molds were warmed with cold tap water to defrost the samples and opened under ethanol to prevent gel rapid swelling. Then, the prepared gels were immediately placed into deionized water. The water was changed frequently for several days until it reached the conductivity of pure water.

### Scanning electron microscopy (SEM)

SEM measurements were obtained using a JSM-6010LA scanning electron microscope (JEOL Ltd., Tokyo, Japan). Samples were freeze-dried, sputter-coated with a gold/palladium mixture using an E-1010 ion sputter system (HITACHI, Hitachi, Japan) and analyzed at an acceleration voltage of 20 kV using a BSE detector.

### Fluorescence observation of hydrogels

The porous hydrogel was equilibrated in 0.1 mM fluorescein isothiocyanate (FITC) solution to stain the gel matrix. Then, the FITC solution in the pores was squeezed out in a water bath for 1 minute. 2D and 3D fluorescence imaging was carried out by using an all-in-one fluorescence microscope (BZ-X700, KEYENCE, Osaka, Japan).

### Measurement of the mechanical and physical properties of the C1A1 porous hydrogel

(i) The stiffness of the C1A1 porous hydrogel was measured as the Young’s modulus under compression using Tensilon RCT 1310A (ORIENTEC CORPORATION, Tokyo, Japan), as described previously^34^. Briefly, a 100 N load cell was used to detect the stress response of the sample to the applied strain. Disc samples 10 mm in diameter were obtained from the gel sheets. The compression velocity for all samples was set to a constant 10% strain per minute (~0.220-0.365 mm/min depending on the sample thickness, which is determined by the gel composition and the degree of swelling). Young’s modulus was calculated from the slope of the beginning of the compressive curve.

(ii) The surface electric charge of the hydrogel was measured as the zeta potential by using a submicron particle size analyzer (Delsa Nano HC, BECKMAN COULTER, Brea, LA, USA) as described previously^34^. Briefly, for detection of the potential, a standard particle suspension (500–600 nm in diameter; Otsuka Electronics, Osaka, Japan) consisting of a polystyrene-latex core with a hydroxypropyl cellulose shell was diluted 300 times with a 10 mM sodium chloride (NaCl) aqueous solution. A specimen sheet presoaked in 10 mM NaCl aq was placed on a quartz block cell with a liquid-flow passage, and the standard particle-dispersed solution was used to fill the passage. The zeta potential was evaluated by laser-Doppler electrophoretic light scattering (ELS) (n = 4).

(iii) The surface electric charge of the pores was measured as the Donnan potential, as described previously^35^. Briefly, the C1A1 porous hydrogel was rinsed in a distilled water bath to remove unreacted reagents until the electroconductivity of the bath water at room temperature, which was monitored with a conductance meter (FiveEasy F30, METTLER-TOLEDO, Columbus, OH, USA), reached 0.5 μS/cm, comparable to that of pure water. Then, the gel was equilibrated in 10^-5^ M NaCl solution. The Donnan potential of the gel as a function of bulk depth was measured by a self-developed potential analyzer constructed from an oscilloscope (HDO6034, TELEDYNE LECROY, Chestnut Ridge, NY, USA), amplifier (8700 CELL EXPLORER, DAGAN, Minneapolis, MN, USA) and manipulator (DMA-1511, NARISHIGE, Tokyo, Japan) of a submicron electrode. A tiny electrode with a 150-nm tip edge diameter was inserted into the gel bulk stored in 10^-5^ M NaCl aq. at 0.8 μm/s velocity. A carbon electrode was used as the reference.

(iv) Measurement of the weight swelling ratio *W* of the C1A1 porous hydrogel. Gels were cut into smaller pieces (usually 10 mm cylindrical sample) and weighed to obtain their weight in equilibrium swollen state (*m_sw_*) and then freeze-dried and weighed to obtain their dry weight (*m_d_*). Freeze-dried samples were soaked in cyclohexane and quickly weighed to obtain the weight of samples in nonsolvent (*m_sw,CH_*). The equilibrium water regain (*W*) and dry samples were calculated according to the following equations^36^:

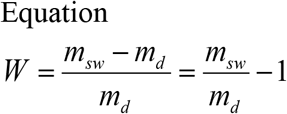

### Preparation of neural stem cells

Neural stem cells (NSCs) were obtained according to a method described previously^37,38^. Briefly, the basal ganglia of mouse brains (Jcl:ICR mouse fetuses, CLEA Japan Co., Ltd., Tokyo, Japan) at embryonic day 14 were fractured by pipetting, and the suspension was disseminated on a culture dish coated with 15 μg/ml poly-L-ornithine (Sigma-Aldrich) and 1 μg/ml human fibronectin (Sigma-Aldrich) (O/F-coated dish). NSCs were maintained in serum-free DMEM and Nutrient Mixture F-12 (DMEM/F-12) (Thermo Fisher Scientific, Waltham, MA, USA) containing 1% N2 supplement (WAKO, Tokyo, Japan). Ten nanograms/milliliter basic fibroblast growth factor (bFGF) (PeproTech, Rocky Hill, CT, USA) was added to the medium every day. After 5 days, cells that reached a proliferative state were used for all experiments.

#### Neutralizing antibody assay for integrin β1

NSCs dissociated in suspension at a concentration of 1×10^5^/ml were incubated with neutralizing antibodies against integrin β1 (AIIB2, DSHB, Iowa City, IA, USA) at concentrations of 0.4 and 2 μg/ml for 15 minutes at room temperature, and the treated NSCs were replated on O/F-coated PS dishes or C1A1 hydrogels. After 1 hour, the cells were fixed with 3% paraformaldehyde (PFA). The numbers of adherent cells were counted by using a phase contrast microscope with a 20× objective lens, and the average of five images was determined. The experiments were repeated four times.

### Immunofluorescence analysis

Cultured cells were fixed with 4% PFA in PBS and permeabilized with 0.2% Triton X-100 in PBS. After blocking with 1% BSA in PBS for 30 min, immunofluorescence was performed as described elsewhere. Briefly, cells or brain tissue sections were incubated with primary antibodies (working dilutions are listed in Table 2) overnight, followed by incubation with secondary antibodies for 1 hour by using an Alexa Fluor-488 anti-mouse IgG antibody (Thermo Fisher Scientific, 1:250), Alexa Fluor-488 anti-rat IgG antibody (Thermo Fisher Scientific, 1:250), Alexa Fluor-594 anti-rabbit IgG antibody (Thermo Fisher Scientific, 1:250), and Alexa Fluor-594 anti-rat IgG antibody (Thermo Fisher Scientific, 1:250). DAPI (Thermo Fisher Scientific) was used for nuclear staining. Fluorescence images were obtained by using a fluorescence microscope (BZ-9000, KEYENCE) and confocal laser scanning microscope (FV 3000, OLYMPUS Co., Tokyo, Japan). Mouse brains were removed after reflux fixation, immersed in 30% sucrose solution overnight, and then frozen at −80°C. Sections of brain tissue (20 μm thick) on glass slides were blocked with 3% bovine serum albumin (BSA) for 1 hour. Immunostaining was performed as described above.

### Establishment of a traumatic brain injury model and implantation of the C1A1 porous hydrogel

A traumatic brain injury model was established by using seven-week-old C57BL/6JJcl or NOD/Shi Jic-scid mice (Table 1). Mice were anesthetized via inhalation with isoflurane and fixed in the cranial position with a specialized device. The skull was exposed, the periosteum was removed, and a circle of thin-layered protective substance with a diameter of 3-4 mm was created on the skull by using the UV light-cured resins GLUMA® Self Etch (KULZER GmbH., Hanau, Germany) and LuxaFlow Star (DMG GmbH, Hamburg, Germany). Subsequently, along with the inner edge of the circle, a hole 3-4 mm in diameter was drilled out. Then, the meninges were removed to expose the cerebral cortex, and cylinder-shaped defects of brain tissues (1 mm in diameter and 1 mm in depth) were generated by aspiration of the brain parenchyma by using a gel loading tip approximately 0.5 μm in diameter. After the bleeding was controlled and the clot was removed with a sterile cotton swab, a C1A1 porous hydrogel with a size of 1 μm square was implanted into the brain tissue defect. Then, the skull hale was covered with a circular cover glass 4.2 mm in diameter (MATSUNAMI GLASS Co., Ltd., Osaka, Japan) and bonded using dental UV-cured resin (Ionosit-Baseliner, DMG GmbH).

**Table 1.**

| Neural stem cells (-) |  |  | Neural stem cells (+) |  |  |  |
| --- | --- | --- | --- | --- | --- | --- |
| conditions after aspiration removal | Day |  | conditions after aspiration removal | Day |  |  |
|  | sacrificed | number |  | NSCs injection | sacrificed | number |
| C1A1 porous hydrogel | 1 | 5 | C1A1 porous hydrogel | 23 | 63 | 5 |
|  | 21 | 5 | Aspiration only | 22 | 64 | 4 |
|  | 56 | 5 | Matrigel | 0 | 35 | 5 |
|  | 200, 294 | 1 of each | C1A1 cryogel (NOD/Shi Jic-scid) | 25 | 61 | 3 |
| Aspiration only | 1 | 5 |  | 20 | 84 | 2 |
|  | 21 | 5 |  |  |  |  |
|  | 56 | 5 |  |  |  |  |

**Table 2.**
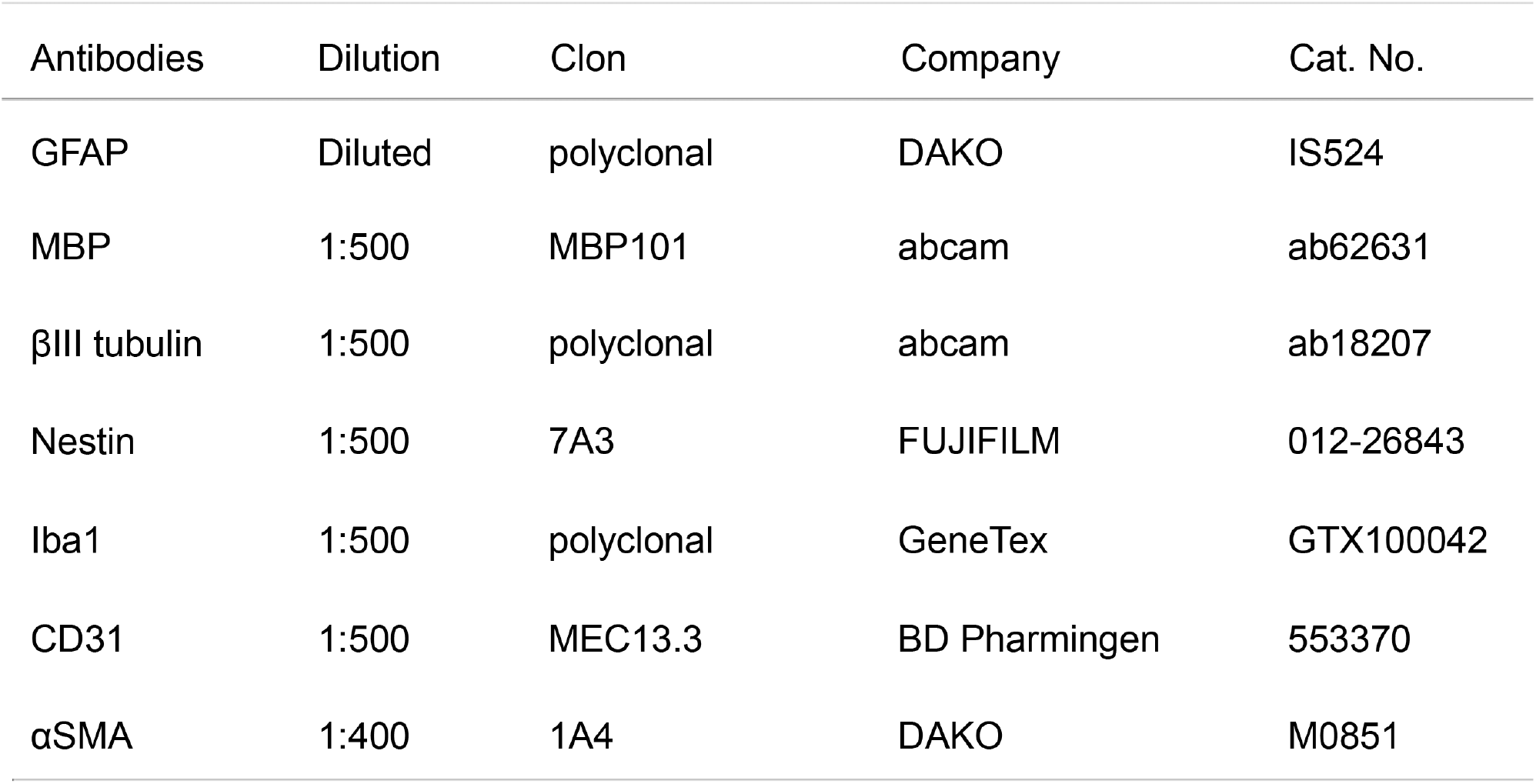

### *In vivo* live imaging

Two-photon fluorescence images were obtained by two-photon laser microscopy customized for *in vivo* imaging (A1R-MP+, Nikon, Tokyo, Japan) with a Nikon Apo LWD 25 ×/1.10 NA (~500 × ~500 μm scanning area) water-immersion objective lens^39^. The correction collar of the objective lens was adjusted by rotating to 0.17. A Ti:sapphire laser (MaiTai HP, Spectra-Physics) was employed as an excitation laser light source at a 920 nm wavelength. Image stacks were acquired with z-steps of 1 μm and 512 × 512 pixels. All fluorescence signals shorter than 690 nm wavelength were detected by non-descanned detectors (NDDs) equipped with GaAsP PMTs. The images were obtained from mice under anesthesia.

### Gene transduction with a lentiviral vector

The lentiviral vector pCSII-EF-IRES-EGFP was used to generate EGFP-expressing NSCs. The lentiviral vector was cotransfected into HEK293T cells (RCB2202, RIKEN, Japan) with psPAX2 and pCMV-VSV-G-RSV-Rev using Polyethylenimine Max (Polyscience Inc., Warrington, PA, USA). Culture supernatants containing the virus were collected at 48 hours after transfection. For infection of NSCs, the lentivirus supernatants were concentrated using 4× polyethylene glycol 6000 (Tokyo Chemical Industry Co., Ltd., Tokyo, Japan) containing 0.4 mM NaCl. NSCs were infected with the lentiviruses in the presence of 10 μg/ml polybrene.

### Cell transplantation

The cranial window was carefully removed to expose the implanted C1A1 porous hydrogel. GFP-labeled NSCs were suspended in PBS at a concentration of 5×10^7^ cells/ml. A Hamilton syringe filled with 2 μl of cell suspension with a 33 G needle was used, and fluids containing a total of 100,000 cells were injected into the implanted hydrogel over 10 minutes. The timing of cell transplantation is shown in Table 1. Cell transplantation was performed at 3 weeks after gel implantation. As a negative control, cell transplantation without a scaffold was performed at 3 weeks after brain defect formation. As another control, cells were transplanted with Matrigel immediately after brain defect formation, with a 30-minute window until the Matrigel was confirmed to be in the solid phase. After completion of these procedures, skin sutures were placed to cover the graft site in all groups. Samples were collected at each time point (Table 1).

#### Image processing

All images were processed with Fiji^40^. The defect area was drawn as an ROI from the cortical surface of the telencephalon to the hippocampus, and the area of the ROI region was measured with Fiji. The total number of cells present in the C1A1 porous hydrogel was measured using DAPI-stained images. The numbers of cells positive cells for each antibody was determined. Glial scarring was assessed by the fluorescence intensity of GFAP per unit area. The regions located 0-20, 20-70, 70-120, and 120-170 μm away from the edge of the hydrogel were delineated as ROIs, and the total fluorescence intensity of GFAP in each region was measured^41^. The viability of the transplanted cells was assessed by the GFP-positive area. The fluorescence images of GFP obtained from three sections in each case were converted to 8-bit, and the GFP-positive areas were masked by the same threshold to measure the area.

#### Statistical analysis

All graphical data are presented as the mean ± s.d. using DataGraph (version 4.6.1, Visual Data Tools, Inc, Chapel Hill, NC, USA). Student’s *t* test was used for the analysis of significant differences; *P* < 0.05 was considered significant, *P* < 0.01, and *P* < 0.001 was considered highly significant. Microsoft® Excel for Mac (version 16.46) was used for all statistical analyses.

## Supplementary Materials

Supplementary movie 1

Three-dimensional image of the C1A1 porous hydrogel.

Supplementary movie 2

*In vivo* live imaging of vascularization in the C1A1 porous hydrogel.

Supplementary movie 3

Three-dimensional culture of neural stem cells using C1A1 porous hydrogel. Red for βIII tubulin and green for myelin basic protein.

Supplementary movie 4

Three-dimensional culture of neural stem cells using C1A1 porous hydrogel. Red for βIII tubulin and green for myelin basic protein. Blue indicates nuclei visualized by DAPI staining.

## Acknowledgments

This work is supported by the Global Center for Soft Matter (a project of the Global Institution for Collaborative Research and Education at Hokkaido University). The Institute for Chemical Reaction Design and Discovery (ICReDD) was established by the World Premier International Research Center Initiative (WPI), MEXT, Japan. The IVIS used in this study was supported by the Global Center for Biomedical Science and Engineering, Faculty of Medicine, Hokkaido University.

## Funding

The Ministry of Education, Culture, Sports, Science and Technology (MEXT) of Japan, Grantin-Aid for Scientific Research: (19K20656 to Sat.T., 19H01171 to Shin.T., 20H05669 to T.N., 21H03802 to M.T., and 21J14773 to T.T.)

Japan Agency for Medical Research and Development (AMED): (20cm0106571h0001, 21cm0106571h0002 to S.T., and Brain/MINDS JP21dm0207078 to T.N.).

## Author contributions

Sat.T. conceived the idea and performed all experiments. Y.E supported the experimental procedures. S.S. generated the charged hydrogels. T.S. established the porous hydrogels and performed scanning electron microscopy. Taka.N., H.K., and Taku.N. measured the physical properties of all gels. J.P.G. organized the projects related to hydrogels. A.K. generated the GFP-positive neural stem cells. T.T., K.Y. performed *in vivo* live imaging supervised by T.N. using a two-photon microscope. M.T, M.I. and Z.T. supported the preparation of the manuscript. Shin.T. designed the entire study.

## Data and materials availability

All data are available in the main text or the supplementary materials.

---

**Supplementary Figure 1.**
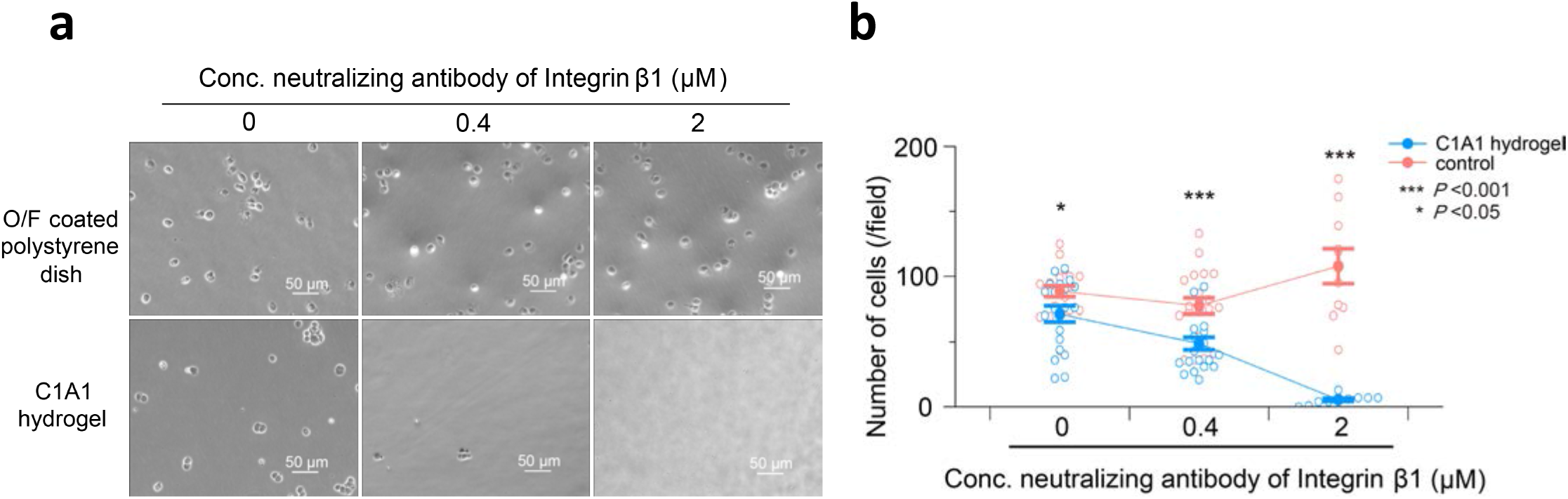
A neutralizing antibody against integrin β1 suppressed NSC adhesion to the C1A1 hydrogel. **a** Phase contrast images of NSCs on the C1A1 hydrogel (lower panels) or poly-L-ornithine/fibronectin-coated polystyrene dishes as controls (upper panels). NSCs treated with the indicated concentrations of integrin β1 neutralizing antibodies were seeded onto C1A1 hydrogels or PS dishes, and after 1 hour, photographs were obtained. **b** The numbers of adherent cells were counted in the presence of the indicated concentration of neutralizing antibody and displayed as plots by using the mean ± SEM. *\*\*\*P* < 0.001. *\*P* < 0.05.

**Supplementary Figure 2.**
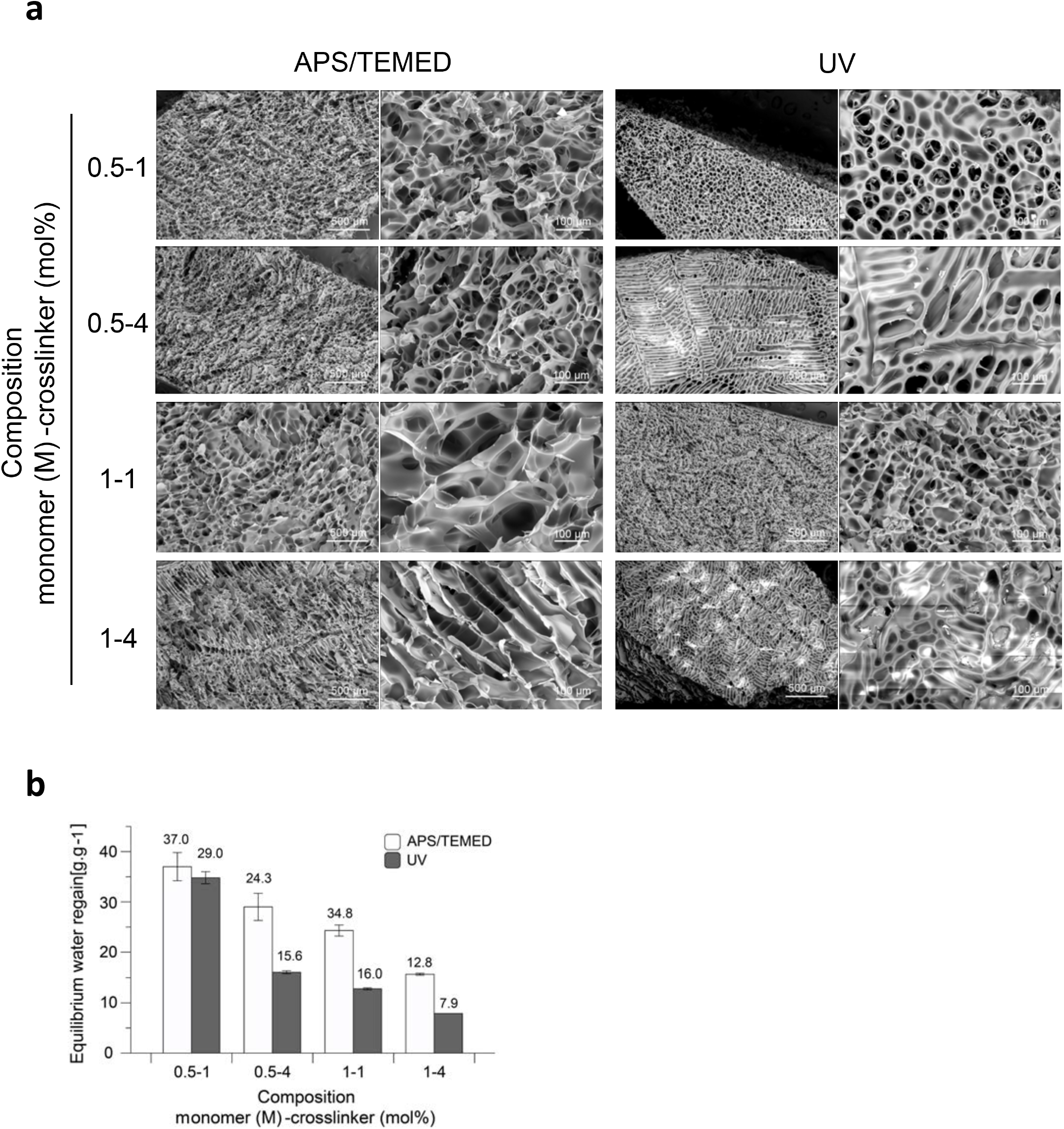
Engineering of C1A1 porous hydrogels with different physical properties. C1A1 porous hydrogels were engineered by using four different combinations of monomer and crosslinker concentrations and two polymerization methods: APS/TEMED or UV irradiation. **a** Scanning electron microscopy images are shown at lower (white bars indicate 500 μm) and higher (white bars indicate 100 μm) magnifications. **b** Value of equilibrium water regain for C1A1 porous hydrogels under each condition are displayed as a bar graph using the mean ±SEM. Open bars and closed bars indicate APS/TEMED and UV polymerization, respectively.

**Supplementary Figure 3.**
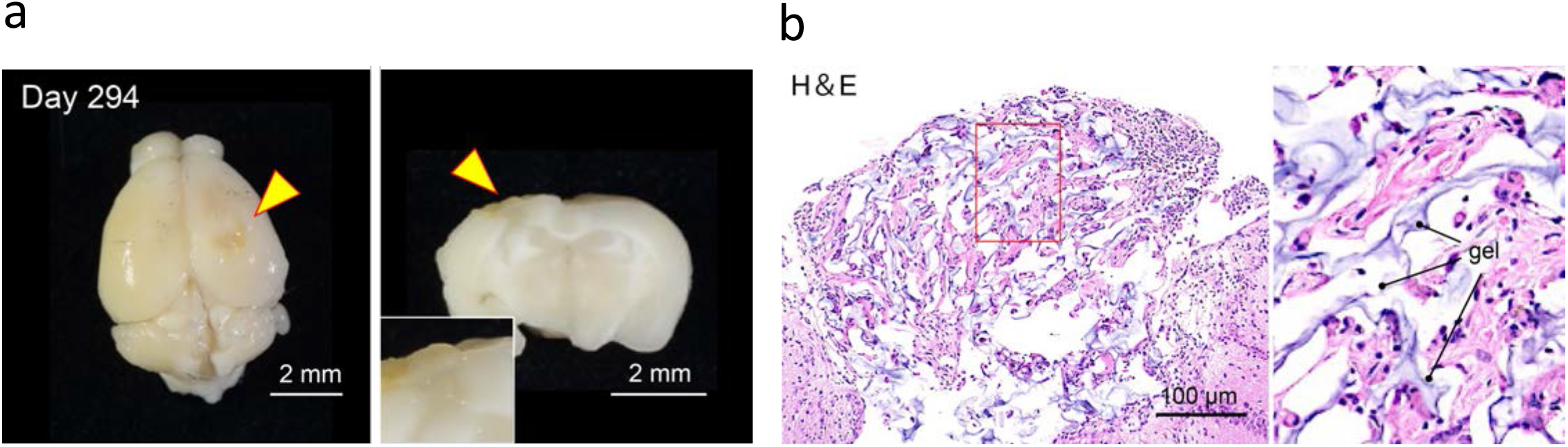
Long-term analysis of C1A1 porous hydrogels implanted into mouse brains. **a** Macroscopic photographs of the C1A1 porous hydrogel implanted into mouse brains on day 294. Yellow arrowheads indicate hydrogels. Whole brain (left) and coronal section (right) of the implanted hydrogel at a higher magnification (inset of right). **b** H&E image of the transplant site. The porous structure of the C1A1 porous hydrogel is shown in light purple.

**Supplementary Figure 4.**
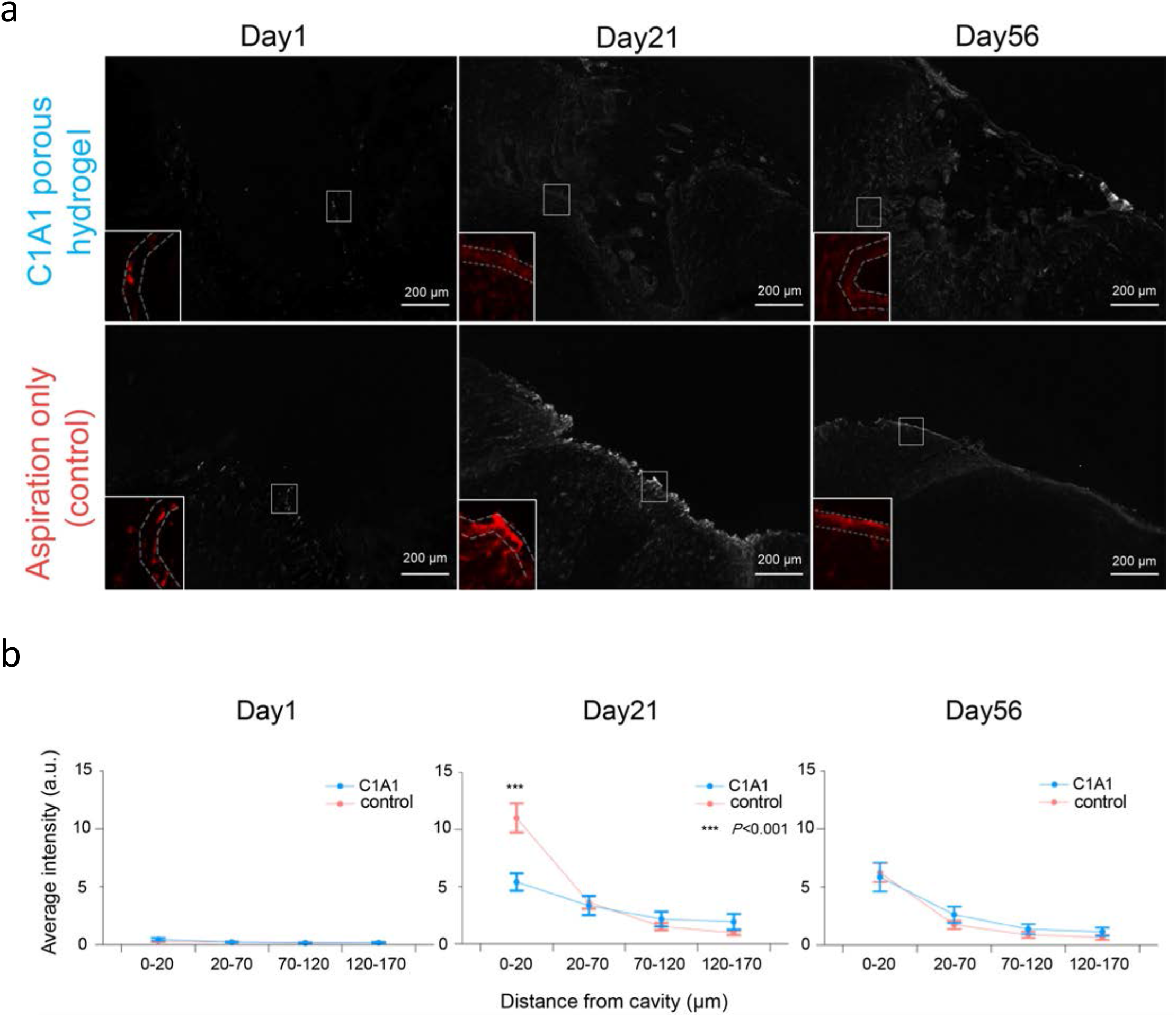
Analysis of GFAP-positive cells at the border of brain defects. **a** Immunofluorescence images of GFAP-positive cells at the border of brain defects with (upper panels) or without (lower panels) implantation of the C1A1 porous hydrogel on days 1, 21, and 56. White bars indicate 200 μm. The insets show a higher magnification. (b) The fluorescence intensities of GFAP were measured and displayed as the mean ± SEM. The immunofluorescence staining images were converted to 8-bit format, and the fluorescence intensities in the regions 0-20 μm, 20-70 μm, 70-120 μm, and 120-170 μm from the boundary were measured. \*\*\**P* < 0.001.

